# Alone Again: Altered Activation in the Observation Execution System during Synchronization in High Loneliness Individuals

**DOI:** 10.1101/2021.10.17.464634

**Authors:** Nira Saporta, Dirk Scheele, Jana Lieberz, Michael Nevat, Alisa Kanterman, René Hurlemann, Simone G. Shamay-Tsoory

## Abstract

Lonely people evaluate social exchanges negatively and display difficulties in interactions. Interpersonal synchronization is crucial for achieving positive interactions, promoting affinity, closeness, and satisfaction. However, little is known about lonely individuals’ ability to synchronize and about their brain activity while synchronizing. Following the screening of 303 participants we recruited 32 low and 32 high loneliness participants. They were scanned while engaged in movement synchronization, using a novel dyadic interaction paradigm. Results showed that high loneliness individuals exhibited a reduced ability to adapt their movement to their partner’s movement. Intriguingly, during movement adaptation periods, high loneliness individuals showed increased activation in the observation-execution (OE) system, specifically in the inferior frontal gyrus and the inferior parietal lobule. They did not show increased activation in the dorsomedial prefrontal cortex, which in the context of synchronization was suggested to be related to gap-monitoring. Based on these findings, we propose a model according to which lonely people may require stronger activation of their OE system for alignment, to compensate for some deficiency in their synchronization ability. Despite this hyper-activation, they still suffer from reduced synchronization capacity. Consequently, synchronization may be a relevant intervention area for the amelioration of chronic loneliness.

## 2 Introduction

Loneliness is a subjective experience of social isolation (Weiss, 1973), perceiving one’s relations as lacking (Perlman & Peplau, 1981). Loneliness is highly prevalent (Barreto et al., 2021; Beutel et al., 2017; Victor & Yang, 2012; Wilson & Moulton, 2010) and has gained public and academic attention as it was shown to harm mental and physical health (Cacioppo et al., 2010; Holwerda et al., 2014; Lim et al., 2020; Valtorta et al., 2016).

Lonely people demonstrate deficits that make engaging in meaningful relationships harder. They experience more negative feelings during interactions (Hawkley et al., 2003) and report lower relationship satisfaction, more conflict, and less self-disclosure and closeness (Mund et al., 2020). Lonely people also maintain larger interpersonal distance (Lieberz et al., 2021), even from friends (Saporta et al., 2021). A potential component of the failure to fully engage in interactions may be related to difficulties in synchronization. Interpersonal synchrony is defined as the alignment in time of the movement of interacting individuals (Chartrand & Lakin, 2013). Synchronization widely occurs naturally, and people coordinate their movement despite not being instructed to (Richardson et al., 2005, 2007). It has been suggested that synchronization evolved to provide important adaptive values (Duranton & Gaunet, 2016; Launay et al., 2016), including achieving emotional alignment (Hatfield et al., 1994) and developing social bonds (Atzil et al., 2011, 2014; Atzil & Gendron, 2017; Feldman, 2007). Indeed, it was found to promote increased liking and affiliation (Hove & Risen, 2009; Rabinowitch & Knafo-Noam, 2015), rapport (Vacharkulksemsuk & Fredrickson, 2012), trust (Launay et al., 2013), empathy (Koehne et al., 2016), connection (Marsh et al., 2009), compassion (Valdesolo & DeSteno, 2011), excitement (Noy et al., 2015) and prosocial behavior, even among infants (Cirelli et al., 2014, for a recent review see Hoehl et al., 2021). It was suggested that perceived social bonding is associated with better synchronization capabilities (Cacioppo & Cacioppo, 2012) and lonely people showed impaired spontaneous smile mimicry (Arnold & Winkielman, 2021). However, despite its significance to achieving significant social interaction, little is known about the ability of lonely people to synchronize with interacting partners.

Synchronization involves several neural networks, most famous of which is the Mirror Neuron System (MNS), which includes neurons in the observation-execution (OE) network, activated both by execution of goal-directed actions and by observation of such actions by another (Rizzolatti et al., 2001; Rizzolatti & Craighero, 2004). Two main areas in the OE system are the inferior frontal gyrus (IFG) and the inferior parietal lobule (IPL) (Iacoboni et al., 2001; Lestou et al., 2008; Pilgramm et al., 2009), and both have been found to be involved in synchronization (Cacioppo et al., 2014; Fairhurst et al., 2013; Jasmin et al., 2016; Jiang et al., 2012; Osaka et al., 2015). Notably, a recent brain model suggested that the OE system is one of three core components of social alignment, which mediate all types of synchrony, from movement to emotional and cognitive alignment (Shamay-Tsoory et al., 2019). In addition to the OE system, this model suggested the existence of a gap-monitoring system, which detects the gap between self and others, comprising the dorsomedial prefrontal cortex (dmPFC), the dorsal anterior cingulate cortex (dACC), and the anterior insula (AI). The model also suggested the existence of a reward system, which signals the gap is optimal, comprising the orbitofrontal cortex (OFC), the ventromedial PFC (vmPFC), and ventral striatum (VS) (Shamay-Tsoory et al., 2019).

Intriguingly, some overlap exists between the alignment networks and the brain areas involved in loneliness. Among high loneliness individuals there is a decrease in white matter density in the IFG (Tian et al., 2014) and the bilateral IPL (Nakagawa et al., 2015). Lesions to the right IFG were associated with decreased loneliness scores, suggesting that the activity of this area in intact brains is related to increased loneliness (Cristofori et al., 2019). Moreover, there is also evidence for the involvement of brain regions that are part of the proposed gap-monitoring and reward systems in loneliness. White matter density is lower among high loneliness individuals in the AI and the dmPFC (Nakagawa et al., 2015; Tian et al., 2014) and high loneliness was associated with lesions to the right AI (Cristofori et al., 2019). Lonely individuals showed reduced self-other representational similarity in the medial PFC (Courtney & Meyer, 2020). They also exhibited blunted functional connectivity between the AI and occipitoparietal regions during trust decisions (Lieberz et al., 2021). Low loneliness individuals had altered functionality of the VS when viewing pleasant social pictures (Cacioppo et al., 2009) and pictures of close-others (Inagaki et al., 2015). (see Lam et al., 2021 for a recent review of structural and functional studies of loneliness).

Based on these findings, the current study examined whether high loneliness individuals show impaired interpersonal synchronization during social interactions. The study used an interactive computerized paradigm, which enables neuroimaging acquisition from participants, as they engage in a joint activity that becomes increasingly synchronized (Marton-Alper et al., 2020). During the study, individuals were scanned using functional magnetic resonance imaging (fMRI) while interacting nonverbally by controlling the movement of differently colored circles. The task included three conditions. A random control condition (rand), in which the scanned participant controlled the movement of one circle, and a computer controlled the second circle; a free movement condition (free) in which both participants saw the circles moved by themselves and by the other participant, and were instructed to move freely, and a synchronized movement condition (sync) in which they were asked to coordinate their movement with the other participant. The rand condition was designed to make it impossible to synchronize, as the movement of the computer-controlled circle was fully randomized and therefore completely unpredictable. In the free condition, spontaneous synchronization could occur. The sync condition was expected to yield the highest level of synchronization. In this study, we focused on a measure of following periods. This is an individual measurement of the relative contribution of each of the dyad members to the achieved synchronization. Based on the host of social impairments experienced by lonely people, it was hypothesized that, as compared to low loneliness participants, high loneliness participants would show diminished following periods, indicating that they contribute less to the achieved synchronization. This was hypothesized to occur in both the free and the sync conditions.

From a neural perspective, the study first aimed to seek support for the interpersonal synchronization neural model. As such, it was hypothesized that during synchronized movement we would see the involvement of the OE system, focusing primarily on the IFG and IPL, as well as the gap-monitoring system (dmPFC, dACC, and AI) and the reward system (VS, vmPFC, and OFC). These regions were expected to be involved in both spontaneous (free movement) and intentional synchronization. Second, as there is some evidence that these areas may be structurally or functionally different among lonely individuals, we hypothesized that differences between high loneliness and low loneliness individuals would also be found in these regions of interest during following periods.

## 3 Materials and Methods

### 3.1 Participants

303 participants were recruited using social media advertisements. Respondents were screened for the following criteria: (i) fluency in Hebrew; (ii) right-hand dominance; (iii) no medication use (except for oral contraceptives); (iv) no history of neurological disorders or psychiatric problems; (v) no conditions that prevented scanning (e.g., a pacemaker, claustrophobia); (vi) normal or corrected-to-normal vision, including no color blindness. In addition, all participants filled the UCLA loneliness questionnaire (Russell, 1996), see details below. The mean UCLA score in the large sample was 40.809 (SD=10.071), median score=39, mode=36. This median score was in accordance was previous findings on similar populations (Russell, 1996). Sixty-eight healthy participants were recruited out of the participants that were screened. Since the study aimed to compare low and high loneliness individuals, half of the group that was recruited had a loneliness score that was higher than the mean score in the larger sample (≥41) and the other half had a loneliness score that was lower than the mean score in the larger sample (<41). Participants were assigned to same-gender dyads. During data analysis, one participant was excluded due to a neurological finding in the anatomical scan. Another participant was excluded since there was an unexplained scan artifact. Two participants were excluded due to excessive head movement during scanning (>2.5mm/°). Therefore, the analyzed sample included 64 participants (45 females, age 18-35, mean age=25.41, SD=4.20). The study was approved by the ethics committee of Tel Aviv University and the institutional review board at the Sheba Tel Hashomer medical center and was conducted in accordance with the latest revision of the Declaration of Helsinki. Participants provided written informed consent to participate in the study.

### 3.2 Experimental Procedure

Each dyad was invited to the center at the same time. It was confirmed that there was no prior acquaintance between them. After joint debriefing, one participant entered the scanner, and the other participant went into a room adjacent to the fMRI scanner control room. Both participants completed the synchronization task (see below), after which the participant in the scanner remained for an anatomical scan. Subsequently, participants switched places and repeated the task. Monetary compensation was provided for participation.

### 3.3 Measures

#### 3.3.1 Synchronization Task

To measure real-time synchronization among interacting participants, the study used a computer-based movement synchronization multi-agent paradigm (Marton-Alper et al., 2020). This game allows individuals to interact nonverbally by controlling the movement of circle-shaped figures with different colors. The displays are fully synchronized as the computers are connected via a closed network. During the game, each player faces a screen with a rectangle presented on it. The participants are instructed to imagine that the rectangle represents a room. At the beginning of the game, two circles appear on the screens, and each player is assigned one of them (blue, red). Participants are instructed to imagine that the circle represents them, as they are moving in the room. The participant in the scanner uses the response box, while the participant outside the scanner uses a keyboard to control the movement of the circles.

The task includes three conditions. 1) Rand condition – each participant controls the movement of the circle that was assigned to them. The other circle’s movement is controlled by the computer and is randomized. The participants are aware that the other circle is controlled by a computer. 2) Free condition - each participant controls the movement of the circle that was assigned to them, and the other circle is controlled by the other participant. Participants are aware that the other circle is controlled by the other participant and are instructed to move their circle freely. 3) Sync condition – this condition is similar to the free condition, however the participants are instructed to synchronize their movement to the best of their ability. The order of the conditions was maintained for all participants as was established in a previous study (Marton-Alper et al., 2020) so that instructed synchrony will not affect the emergence of spontaneous synchrony right after it.

Prior to entering the scanner, participants received an explanation about the task and were shown the response box they would be using. Each condition was scanned in a separate run and contained 3 blocks. At the beginning of each block, a fixation point appeared on the screen for 12 seconds, followed by the presentation of an instruction slide (5 seconds), after which the participants performed the task for 45 seconds. After this, participants were given 10 seconds to rate how much they enjoyed the game. Figure 1 presents an illustration of the task design.

**Figure 1:**
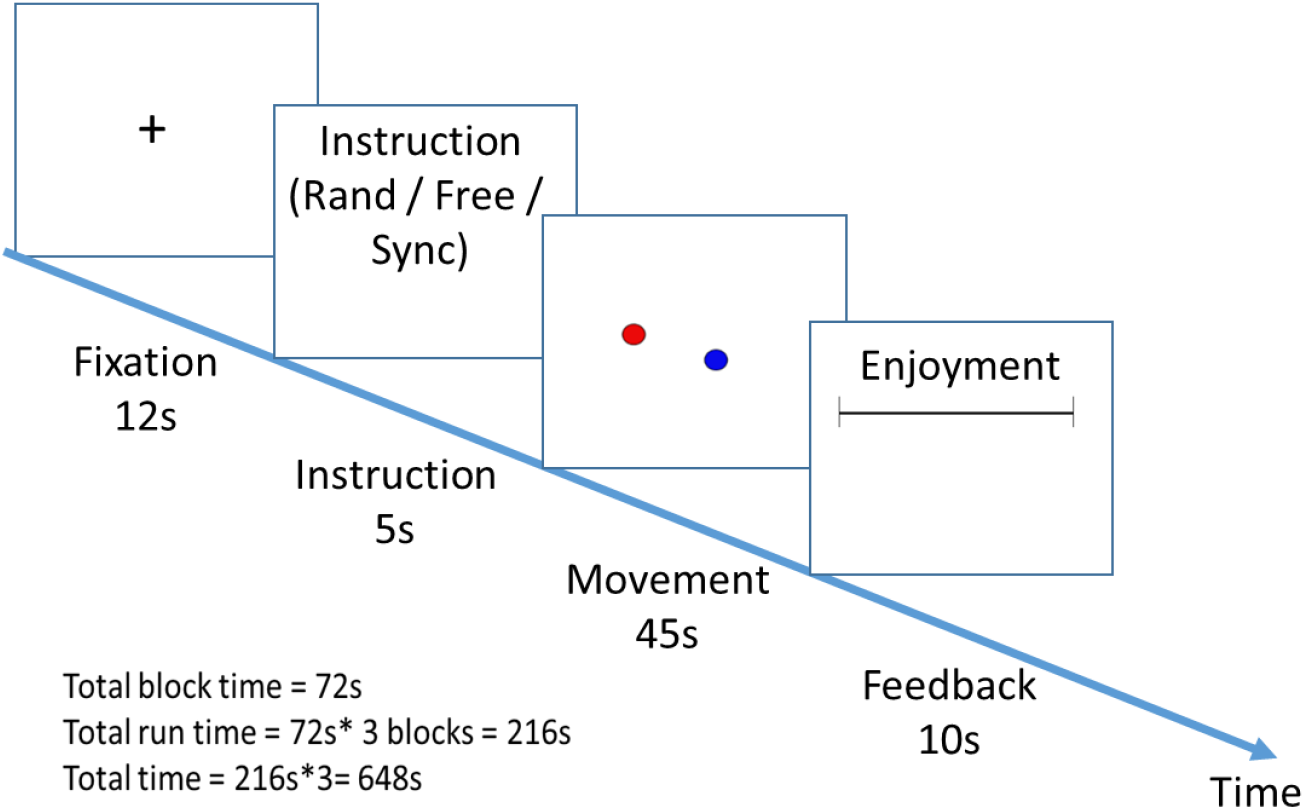
Illustration of the task design

The movement of the circle assigned to the participant outside the scanner was controlled using the 1-4 keys on a keyboard, and the movement of the circle assigned to the participant inside the scanner was controlled using the 4 keys of the response box. Each key represented a direction (left, up, right, down) and combinations of two keys were allowed (e.g., left+up = diagonal left). The speed of movement of each circle was determined by the vector sum of movement in the four major directions, proportional to the duration for which each respective key was pressed. Communication between each client and the server was executed asynchronously at about 5 Hz and post-processing interpolation of all data was conducted at a rate of 5 Hz, such that data for all participants shared matching sample times.

#### 3.3.2 Synchronization Measurements

##### 3.3.2.1 Following Period

To measure the relative contribution of each participant to the achieved synchronization, we examined periods during which a participant actively adapted their movement to that of their partner. As mentioned above, each participant’s location was recorded at a sampling frequency of 5 Hz, yielding 5*45=225 samples per block. From this data we calculated changes in locations between consecutive samples and used these differences to identify the direction in which each participant had moved during each interval between consecutive samples. There are 8 possible movement directions (0°, 45°, 90°, 135°, 180°, 225°, 270°, and 315°). The net change in location between two consecutive samples is a vector sum of the products of directions selected by the participant within that interval, multiplied by their durations. This sum does not necessarily coincide with one of the eight aforementioned directions. Therefore, actual directions were “rounded” to the closest main direction. Periods during which the two dyad members were moving in the same direction were then identified. Within these periods, we projected participants’ locations onto their (common) direction of movement. The person who was “behind” was considered to be the one that was following the person who was “in front”. A following period for participant 1 was defined as the period in which participant 1 was the one following participant 2. A total **following score** was calculated, summing up the following periods per participant and per condition. A higher score would indicate that a participant spent more time actively aligning their movement direction with the movement of the other participant.

##### 3.3.2.2 Zero-lag correlation

To validate the task worked as expected in the scanner, we also analyzed participants’ behavior using a previously created dyad measurement of synchronization, the **zero-lag correlation score** (Marton-Alper et al., 2020). This measurement is based on a directional correlation (Nagy et al., 2010) between the movement of the two participants. Directional correlation is the cosine of the angle between the velocities of each pair of players. The directional correlation between participant i and participant j is given by 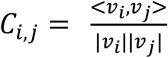, where <v_i_,v_j_> is the inner product of the velocities of the two participants, and |v_i_| and |v_j_| are the magnitudes of the velocities of participant i and participant j, respectively. Higher correlation indicates stronger synchronization. The zero-lag correlations were calculated using sliding windows of 21 samples (i.e., ±2 sec around a given sample). Figure 2 presents an example of the synchronization score calculated over time for one of the dyads.

**Figure 2:**
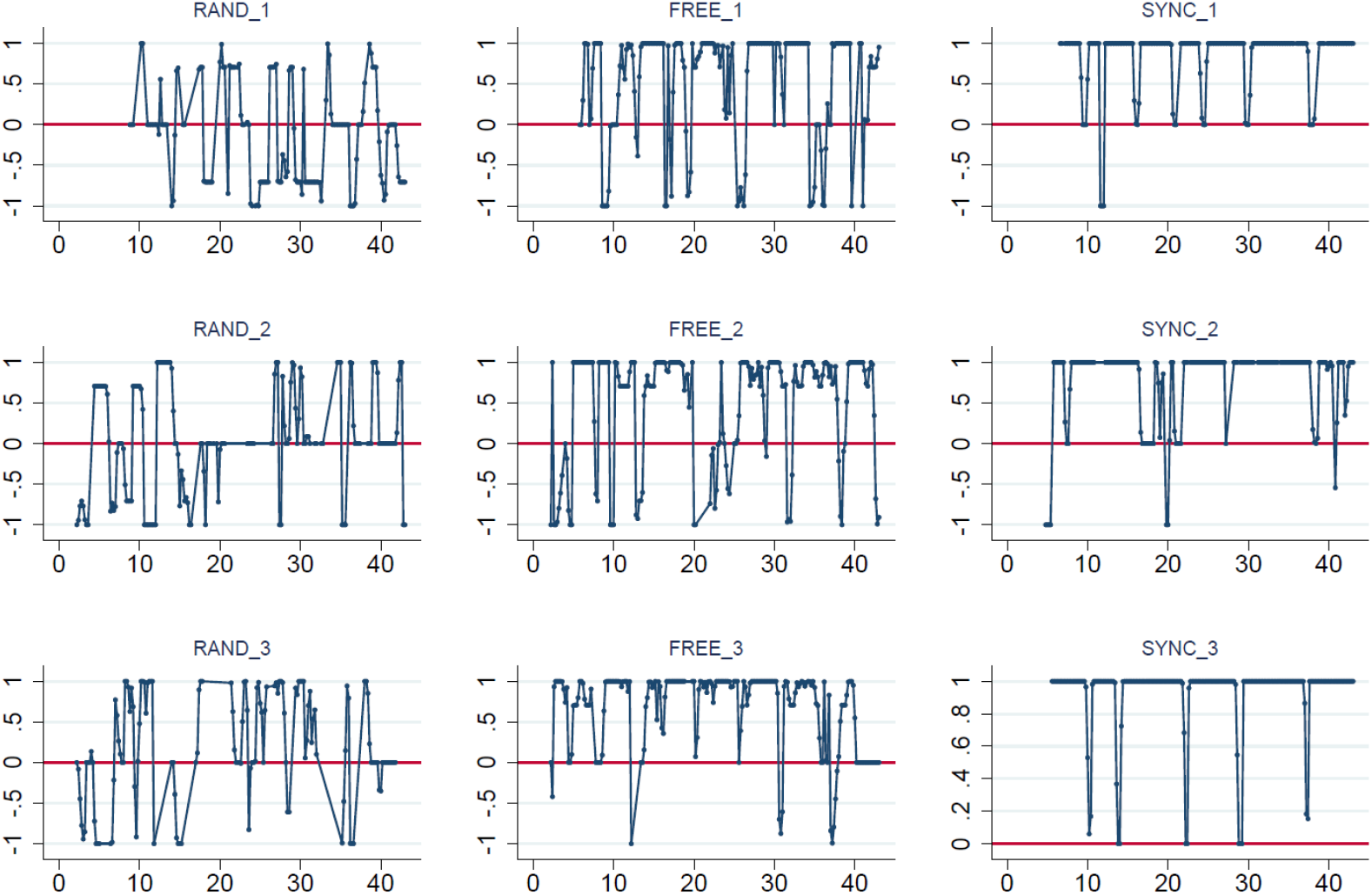
Example of the zero-lag synchronization scores calculated over time for one dyad. The blocks of rand condition (left), free condition (middle) and sync condition (right) are presented. The X axis represents the time in seconds, and the Y axis is the synchronization score

While the zero-lag correlation score is a valuable measurement for the level of synchronization a dyad achieves, it does not allow for a differentiation of the relative contribution of each of the participants to this synchronization. Both members of the dyad will get the same zero-lag correlation score, even though one may have contributed far more to the synchronization, by adjusting his or her movement more. It was especially important to differentiate the individual contribution of the participants since the study aimed to examine individual brain activation. However, since following score is a new measurement, we also behaviorally analyzed zero-lag correlations to assure the task works as expected in the fMRI environment.

#### 3.3.3 Enjoyment Ratings

To further validate the task and the behavioral differences between the three conditions, participants’ enjoyment ratings in each of the blocks were used to calculate an average enjoyment score in each condition for each participant.

#### 3.3.4 Loneliness Measurement

To assess levels of loneliness, participants completed the UCLA loneliness scale version 3 (Russell, 1996). The UCLA loneliness scale was initially developed in 1978 (Russell et al., 1978) and has since been revised twice to improve its validity and reliability. In the current version the respondent is asked to rate the frequency of loneliness-related experiences. Some items refer to negative experiences, for example “How often do you feel left out?” and some items refer to positive experiences, for example “How often do you feel part of a group of friends?”. Each item is rated on a scale of 1 (never) to 4 (often), and after reversing the questions that relate to positive experiences a total loneliness score (20-80) is calculated. The mean score in the UCLA scale in the study was 41.938 (SD=12.952) and the median score was 38. As explained in the participants section, participants filled out the UCLA scale during screening, and then two groups were recruited, based on their loneliness score. Comparing high and low loneliness groups was done in multiple past studies (e.g., Arnold & Winkielman, 2021; Cacioppo et al., 2015; Cacioppo et al., 2016). The mean loneliness scores in the low and high loneliness groups were 30.593 (SD=4.550) and 53.281 (SD=7.385), respectively.

### 3.4 Data Analysis

#### 3.4.1 Behavioral data analysis

Behavioral data were analyzed by calculating mixed-design analyses of variance (ANOVAs), with either following score, zero-lag correlation score or enjoyment score as the dependent variable, condition (rand, free, or sync) as the within-subject repeated measure and loneliness group (high, low) as the between-subject factor. Additional analyses included t-tests and bi-variate Pearson correlations. p-values < 0.05 (two-tailed) were considered significant. Effect sizes were estimated using partial eta squared (ηp2) or Cohen’s d. Cronbach’s Alpha was calculated on the UCLA loneliness scale as a measure of its reliability. All statistical analyses were performed using SPSS 25.0. As each participant performed the synchronization task twice, once outside the scanner and once inside the scanner, scanning order was used as a between-subject control variable to test if it impacted the results.

#### 3.4.2 MRI Data acquisition

MRI was conducted using a 3T Siemens Magnetom Prisma Scanner (Siemens Medical Solutions, Erlangen, Germany) at the Strauss Imaging Center on the campus of Tel Aviv University. Images were acquired using a 64-channel head coil. Every session included 3D-anatomical scanning and functional imaging. Anatomical scans were obtained using a T1-weighted 3D MP2RAGE (TR - 2.53 s; TE - 2.99 ms; flip angle - 7°, 176 sagittal slices; spatial resolution – 1 × 1 × 1mm^3^). During task performance behavioral judgement was collected via a fiber optic response pad (Current Designs, Inc. PA, USA). Functional MRI was acquired by multiband echo planar imaging (mb-EPI) pulse sequence for simultaneous excitation for multiple slices with the following parameters: TR=2s, TE=30ms, band factor =2, Ipat=2, isotropic spatial resolution of 2mm^3^ (no gaps).

#### 3.4.3 MRI Data pre-processing and analysis

FMRI data were pre-processed and analyzed using the Statistical Parametric Mapping toolbox for Matlab (SPM12: Wellcome Trust Center for Neuroimaging, University College London). Pre-processing of functional scans included quality assurance, slice timing correction, realignment, corerigstration, normalization to a standard T1 template (MNI) and smoothing. Head movement was assessed and corrected. 3D statistical parametric maps were calculated separately for each subject using a general linear model (GLM). First level contrasts of interest were calculated (see below), and then we used a one-sample t-test analysis on the second level. All GLM analyses were thresholded at a family wise error (FWE) corrected whole brain p value < 0.05 after an initial cluster-forming height threshold of P < 0.001. We contrasted each of the conditions in which participants had been interacting with each other (sync or free, separately) with the rand condition. Obtained whole brain analyses were masked with the activation map obtained for the sync/free condition minus its baseline (p<0.05), to ensure that the resulting differences were due to activation in that condition rather than deactivation in the rand condition. To determine whether scanning order had impacted the results, we also performed second level analyses in which scanning order was included as a between-subject factor, and a two-sample t-test was conducted to compare the two scanning order groups.

Two types of whole-brain analyses were implemented at the individual level. First, to validate the proposed neural model for interpersonal synchronization and to explore the three conditions of the task, regardless of the actual behavior of the subjects in each task condition, we carried out an initial contrast between the task conditions. We contrasted *individual brain activity throughout the sync condition with the rand condition (total run duration sync>rand)* as the main contrast of interest on the first level and then used a one-sample t-test analysis on the second level. A similar analysis was done for the free condition *(total run duration free>rand)*. The contrasts used the onsets and durations of the task condition, summing up the 3 movement trials of 45 seconds each. Second, to explore brain activation during following periods, as defined in section 3.3.2.1, we contrasted individual brain activity for *following periods in the sync condition > following periods in the rand condition* as the main contrast of interest on the first level. The following periods were significantly longer in the sync condition compared to the rand condition. To assure similar durations were used, and since there was no expectation for intentional following in the rand condition, the onsets and durations of following were duplicated and used in the rand condition as well. A similar analysis was done for the free condition *following periods free condition > following periods rand condition*.

We also conducted an ROI analysis focused on the difference between high and low loneliness participants. Our hypothesis focused on the brain regions suggested to be involved in synchronization (dmPFC, dACC, AI, IFG, IPL, premotor cortex, OFC, vmPFC, VS). Out of those, we identified the ROIs in which activation was confirmed in the whole brain analysis of following periods as described above, as this analysis established their relevance to brain activity during following periods. Anatomical ROIs were then defined using the Automated Anatomical Labeling atlas version 3 (Rolls et al., 2020). Beta values were extracted, and then used to test differences between the high and low loneliness groups using. False discovery rate (FDR) correction was used for multiple comparisons. p values smaller than 0.05 after correction were considered significant.

## 4 Results

### 4.1 Behavioral Data

The reliability of the UCLA Loneliness Scale was excellent (Cronbach’s Alpha=0.957). A mixed-design ANOVA with following score as the dependent variable was employed. The analysis yielded significant main effects of condition [F_(1.148,71.152)_=149.431, p<0.001, η_p_^2^=0.707] and of loneliness [F_(1.148,71.152)_=4.764, p=0.033, η_p_^2^=0.071]. A significant interaction between condition and loneliness was also found [F_(1.148,71.152)_=5.503, p=0.018, η_p_^2^=0.082]. Following score was significantly higher in the sync condition compared with the rand condition [t_(62)_=11.911, p<0.001, Cohen’s d=2.088] and compared with the free condition [t_(62)_=12.300, p<0.001, Cohen’s d=2.117]. The difference between the free and the rand condition was not significant (p=0.696). Follow-up analysis revealed that the high loneliness group showed a lower following score in the sync condition compared with the low loneliness group [t_(62)_= 2.373, p=0.021, Cohen’s d=0.593]. See Figure 3.

**Figure 3:**
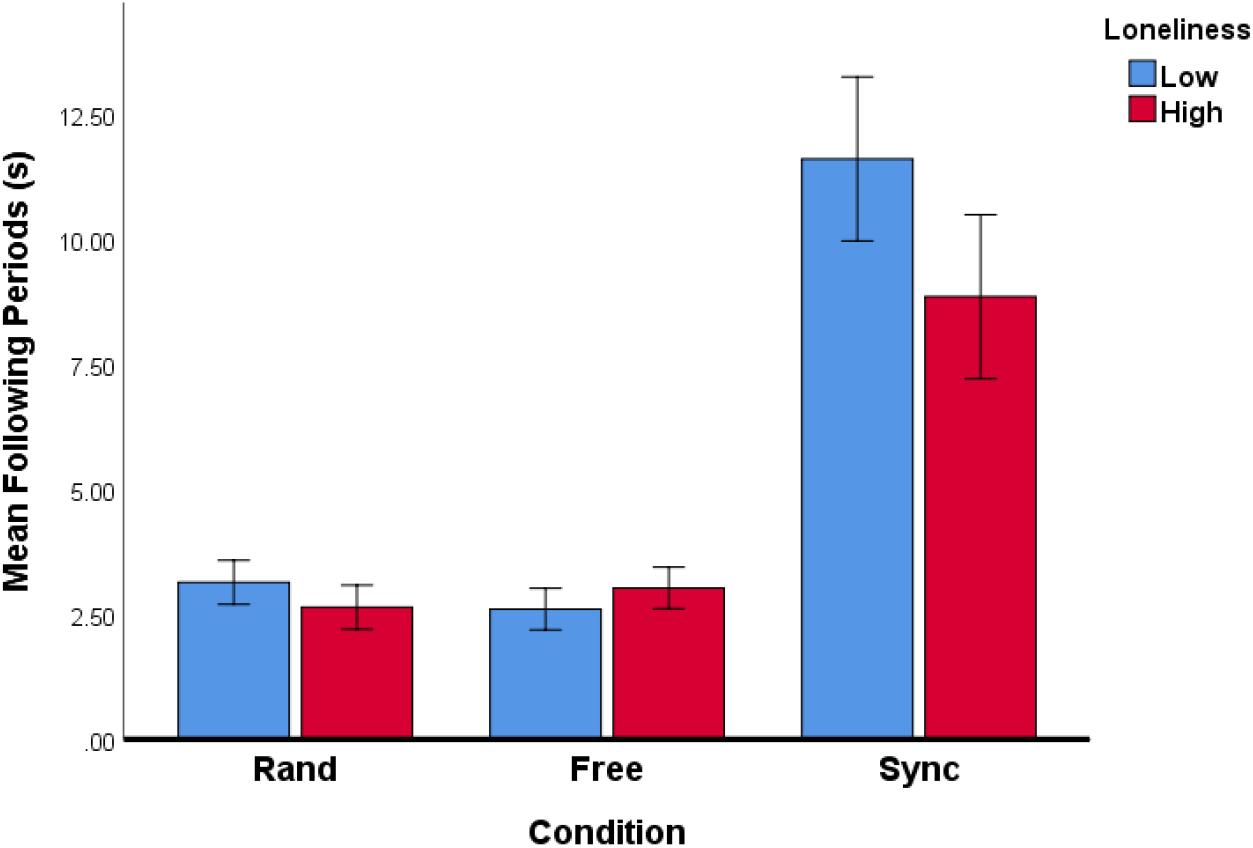
High loneliness group had a lower following score in the sync condition. following score was higher in the sync condition, compared to free and rand condition. Error bars=95% confidence level

A mixed-design ANOVA with zero-lag correlation score as the dependent variable yielded a significant main effect of condition [F_(1.706,105.784)_=343.110, p<0.001, η_p_^2^=0.847]. No other main effects or interactions were significant (p>0.404). As expected, the zero-lag correlation was significantly higher in the sync condition (M=0.522, SD=0.165) compared with the free condition (M=0.011, SD=0.149) [t_(62)_=19.231, p<0.001, Cohen’s d=3.251] and compared with the rand conditions (M=0.008, SD=0.078) [t_(62)_=24.110, p<0.001, Cohen’s d=3.983]. There was no significant difference between the free and rand conditions (p=0.902). When comparing high and low loneliness groups’ zero-lag correlations scores, there was no significant difference in any of the conditions (p≥0.344).

Furthermore, we explored differences in enjoyment between the conditions. A mixed-design ANOVA with the enjoyment score as the dependent variable was employed. The analysis yielded a significant main effect of condition [F_(1.761,109.167)_= 8.856, p<0.001, η_p_^2^=0.125]. No other main effects or interactions were significant (p>0.162). Enjoyment level in the sync condition (M=62.787, SD=19.956) was higher when compared to the rand condition (M=54.037, SD=24.359) [t_(62)_= 3.657, p=0.001, Cohen’s d=0.393]. The enjoyment level in the free condition (M=60.208, SD=22.219) was also higher when compared to the rand condition [t_(62)_=2.803, p=0.007, Cohen’s d=0.265]. The difference between the enjoyment levels in the free condition and the sync condition was not significant (p=0.136). When comparing high and low loneliness groups enjoyment scores, there was no significant difference in any of the conditions (p≥0.173).

To confirm no difference existed between participants who were scanned first and those participants who were scanned second in parameters of age and loneliness, we ran t-tests with UCLA loneliness score and age as the dependent variables, and order as the between-subject factor. These analyses yielded no significant differences between the two groups (p>0.544). When including order as an additional between-subject factor in the mixed-design ANOVAs reported above (with following score or zero-lag correlation score or enjoyment score as the dependent variable), the main effects and/or interactions reported were not impacted by order, and no interactions or main effects with order were found (p> 0.072).

### 4.2 Neuroimaging analysis

Whole brain analysis comparing activity patterns between the total run duration of the sync condition and the rand condition showed that during the sync condition there was increased activation in the right IPL (58, -46, 32), right IFG opercular part (50, 18, 34), left IPL (−56, -58, 28), and dmPFC (18, 60, 24). In addition, there was increased activation in left superior cerebellum (−22, -76, -34) and the middle and superior temporal gyrus*/*STS (46, -24, -6), see Figure 4 and Table 1. A similar whole brain analysis was conducted, comparing activity patterns between the total run duration of the free condition and the rand condition. During the free condition there was increased activation in the right supramarginal gyrus and right IPL (52 -44 24) and the lateral surface of the superior frontal gyrus extending to the mPFC (18, 56, 22). Additional activation was detected in the supplementary motor area (16, -2, 74), see Figure 5 Table 1. We repeated the analyses, including scanning order as a between-subject factor, conducting a two-sample t-test analysis on the second level, and no significant differences in activation were found. These findings show that areas in the OE network and the gap-monitoring network were active throughout both the sync condition and the free condition, with more activation in the sync condition, when compared to the rand condition.

**Table 1:**
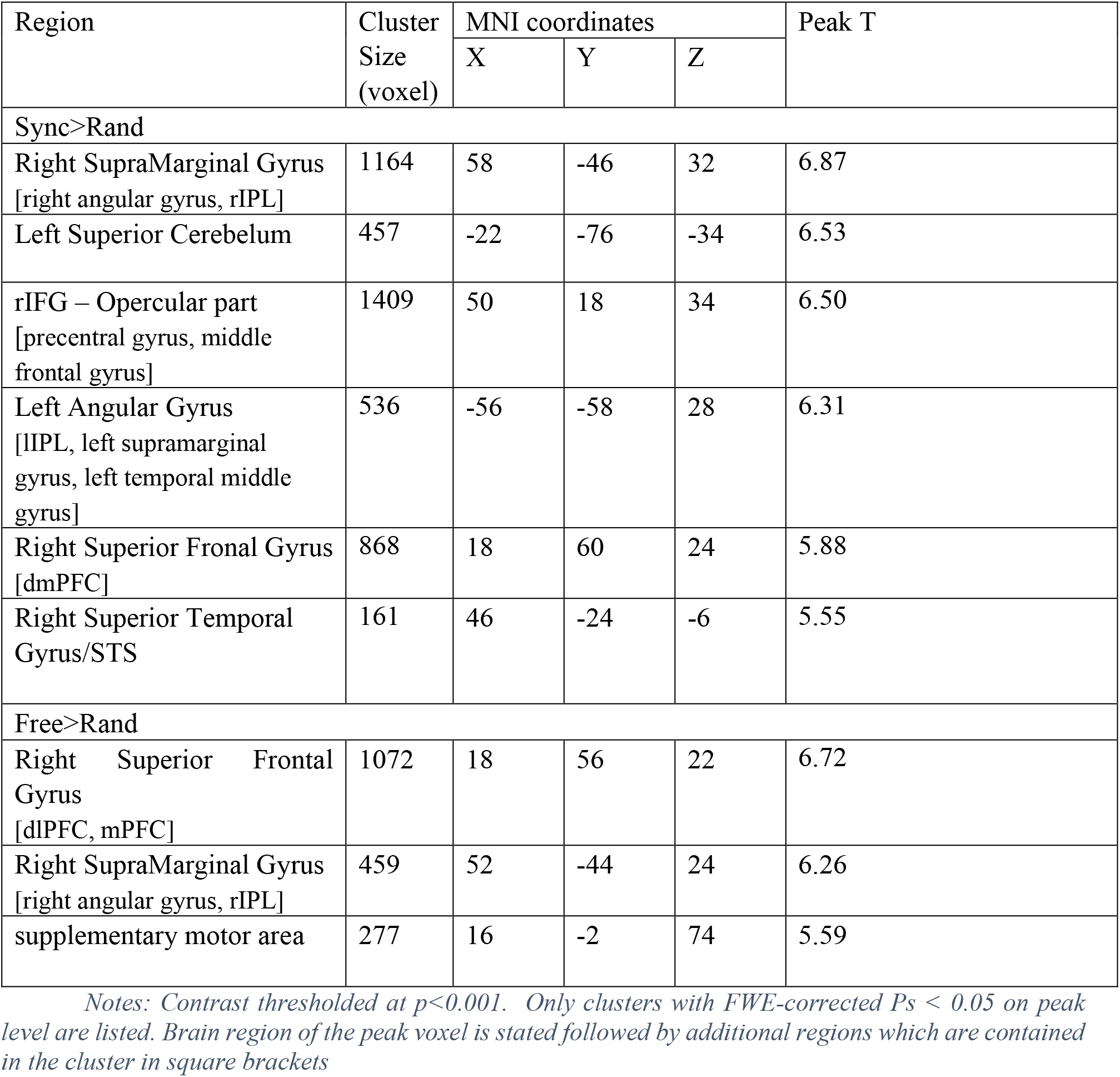
Whole brain analysis sync condition>rand condition throughout the entire condition duration and free movement condition>rand condition throughout the entire condition duration

**Figure 4:**
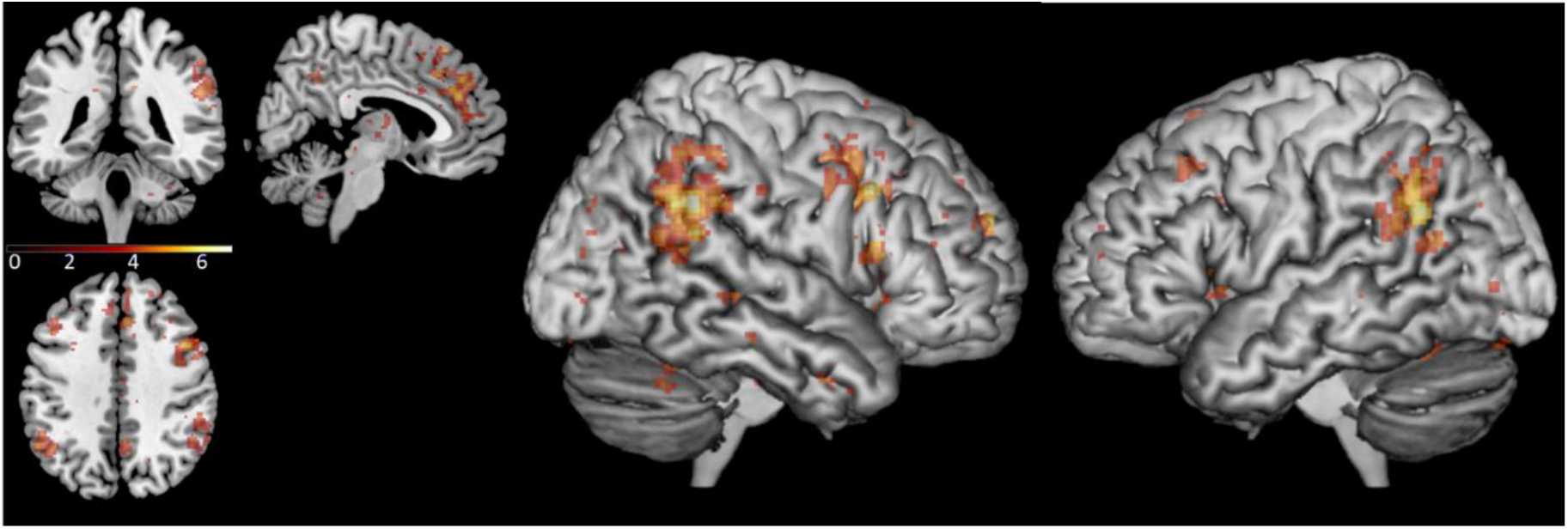
Brain activation in the sync condition>rand condition contrast, during the total run duration, across loneliness groups. Contrast thresholded at p<0.001 for the illustration. MNI coordinates of Axial/Coronal/sagittal view - (5, -40, 40).

**Figure 5:**
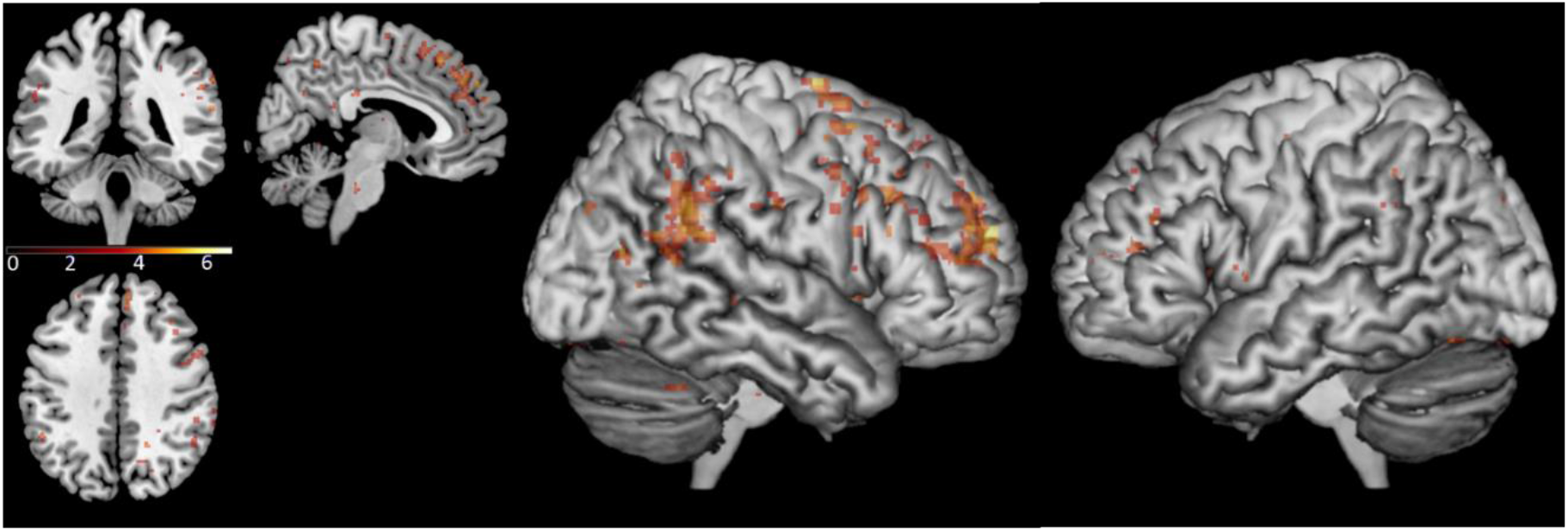
Brain activation in the free condition>rand condition contrast, during the total run duration, across loneliness groups. Contrast thresholded at p<0.001 for the illustration. MNI coordinates of Axial/Coronal/sagittal view - (5, -40, 40)

To analyze the differences in brain activation during periods in which the scanned participant was actively aligning their movement with the other participant (as opposed to the entire run duration of each condition, regardless of the specific behavior of the participant in that time), whole brain analysis comparing activity patterns during following periods in the sync condition (see explanation in section 3.3.2.1) and parallel periods of time during the rand condition. This yielded significant clusters in the right IPL (54, -42, 48), the left IPL (−56, -46, 38), the right IFG opercular part (42, 8, 50), and the dmPFC, extending also into the lateral surface of the superior frontal gyrus (14, 26, 62). In addition, there was increased activation in the superior cerebellum (−18, -78, -26) and the right middle occipital gyrus (38, -84, 22). See Figure 6 and Table 2. A similar analysis of the free condition compared to the rand condition did not yield significant clusters. We repeated the analyses, including scanning order as a between-subject factor and conducting a two-sample t-test analysis on the second level, and no significant differences in activation were found. These findings show that areas in the OE network and the gap-monitoring network were active throughout following periods in the sync condition.

**Table 2:** Whole brain analysis, following periods in the sync condition>rand condition

| Region | Cluster Size<br>(voxel) | MNI coordinates |  |  | Peak T |
| --- | --- | --- | --- | --- | --- |
|  |  | X | Y | Z |  |
| Left Superior Cerebellum | 381 | -18 | -78 | -26 | 7.36 |
| Right SupraMarginal Gyrus<br>[rIPL, right angular gyrus] | 764 | 54 | -42 | 48 | 7.14 |
| Right middle occipital gyrus | 1199 | 38 | -84 | 22 | 6.84 |
| Superior frontal gyrus<br>[dmPFC] | 503 | 14 | 26 | 62 | 6.44 |
| lIPL<br>[left angular gyrus] | 260 | -56 | -46 | 38 | 6.31 |
| rIFG, opercular part<br>[precentral gyrus] | 849 | 42 | 8 | 50 | 5.89 |

Notes: Contrast thresholded at $p < 0.001$ . Only clusters with FWE-corrected $P_s < 0.05$ on peak level are listed. Brain region of the peak voxel is stated followed by additional regions which are contained in the cluster in square brackets

**Figure 6:**
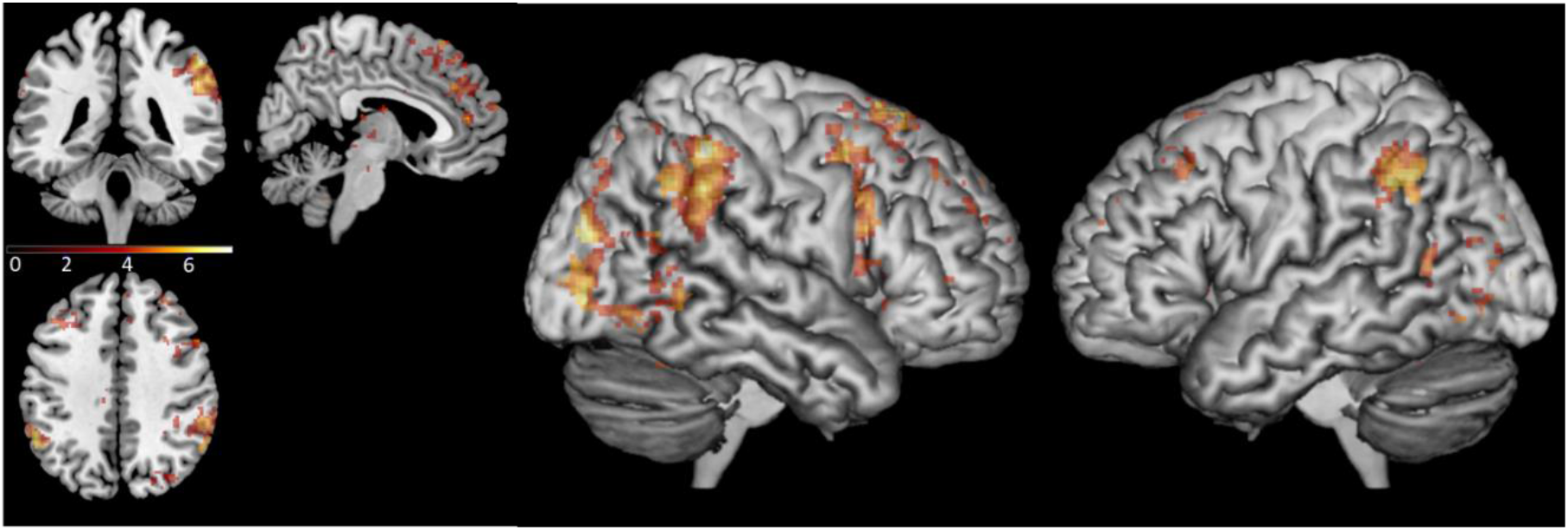
Brain activation in the following periods sync>rand contrast, thresholded at p<0.001 for illustration. MNI coordinates of Axial/Coronal/sagittal view - (5, -40, 40).

As whole brain analysis confirmed activation in the IPL, the IFG, and dmPFC during following periods, beta values were extracted from the sync>rand following periods contrast using the relevant anatomical ROIs. An independent sample t-test analysis revealed a significant difference in activation between the high loneliness group, which had a higher activation in the left IFG (M=0.254, SD=0.375) compared to the low loneliness group (M=0.023, SD=0.349) (t(_62)_=2.547, p=0.013, Cohen’s d=0.637). Similarly, in the right IPL the high loneliness group had a higher activation (M=0.528, SD=0.567) compared to the low loneliness group (M=0.216, SD=0.432) (t_(62)_=2.478, p=0.016, Cohen’s d=0.619). The differences in the dmPFC, lIPL, and the rIFG were not significant (p>0.151). T-tests were FDR corrected for multiple comparisons. See Figure 7 and Table 3 for detailed results.

**Table 3:** Comparing high and low loneliness groups in regions of interest activity during following periods

| ROI | Low loneliness<br>(M±SD) | High loneliness<br>(M±SD) | t-test | FDR<br>corrected p<br>value |
| --- | --- | --- | --- | --- |
| lIFG | 0.023 ± 0.349 | 0.254 ± 0.375 | (t(62)=2.547, p=0.013) | 0.040* |
| rIPL | 0.216 ± 0.432 | 0.528 ± 0.567 | (t(62)= 2.478, p=0.016) | 0.040* |
| rIFG | 0.192 ± 0.329 | 0.332 ± 0.367 | (t(62)=1.609, p=0.113) | 0.188 |
| lIPL | 0.117 ± 0.565 | 0.319 ± 0.589 | (t(62)=1.401, p=0.166) | 0.208 |
| dmPFC | 0.259 ± 0.592 | 0.378 ± 0.488 | (t(62)=0.877, p=0.384) | 0.384 |

Note: \* $p < 0.05$

**Figure 7:**
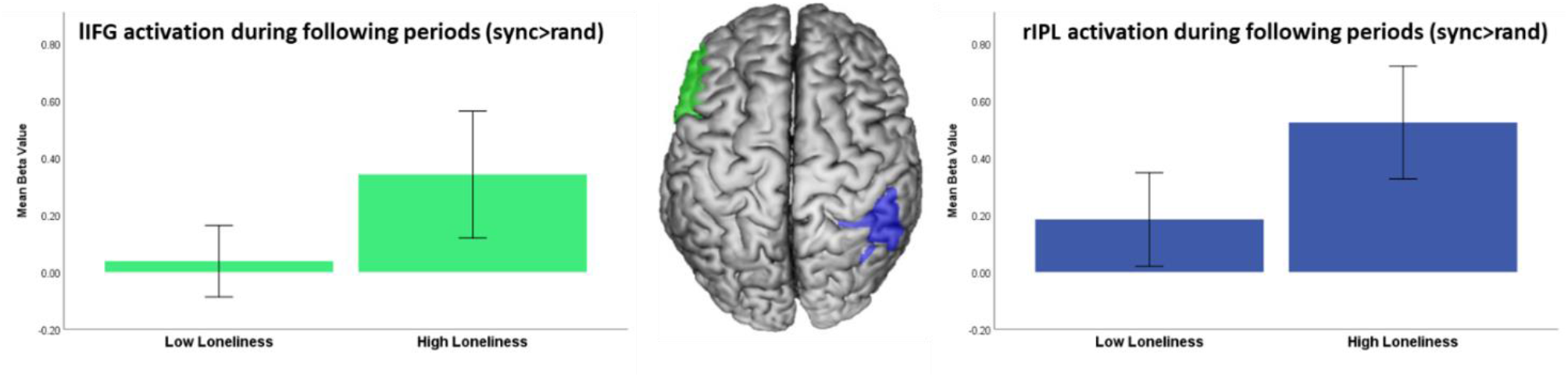
high loneliness group presents a higher activation in the lIFG and rIPL during following periods. Error bars=95% confidence level (cl)

## 5 Discussion

In this study we set out to examine whether lonely individuals have a reduced ability to synchronize with others. Furthermore, we tested a proposed neural model for interpersonal synchronization and explored the neural activation related to synchronization in high and low loneliness individuals. Using a novel computerized fMRI paradigm, we were able to measure neural activity during naturalistic, live interaction of participating dyads.

Our initial hypothesis was confirmed, as the high loneliness group showed a lower level of ability to synchronize, which was reflected by their lower following scores in the sync condition. This observation supports previous findings with regards to the social impairments of lonely individuals (Mund et al., 2020), and sheds additional light on the underlying mechanisms that may inhibit positive social interaction for lonely individuals. If lonely individuals have difficulties in aligning themselves with others, they will most likely miss out on the social benefits of synchronization such as increased connection, engagement, satisfaction, liking and affiliation (Hoehl et al., 2021), which in turn may result in their more negative reports of their interactions.

The results of our study support the model proposed for interpersonal synchrony (Shamay-Tsoory et al., 2019). When examining whole brain activity during synchronization, activations were observed in the IFG and IPL (related to the OE system) as well as in the dmPFC (related to the gap-monitoring system). These findings, which are based on measuring brain activity during a naturalistic and interactive interpersonal synchronization, further strengthen the notion that interpersonal synchronization does not involve only sensorimotor components, but that indeed additional neural networks are recruited. Specifically, our study provides support for the existence of the proposed gap-monitoring system, which assists in obtaining synchronization.

Intriguingly, when examining high and low loneliness individuals engaged in active synchronization using the measurement of following periods, high loneliness was related to increased neural activity in the IPL and the IFG. This suggests that high loneliness individuals need to activate their OE system more when they are asked to synchronize their movement, compared to low loneliness individuals. These hyper-activations may be related to social impairment in loneliness. Considering that the OE system contributes not only to motor alignment, but also to emotional and cognitive alignment (Shamay-Tsoory et al., 2019), this may be relevant to other types of social interaction as well. Similar findings of OE increased activation were reported also for other conditions. For example, Minichino & Cadenhead, (2017) proposed that hyperactive states of the OE may be related to the social deficits in schizophrenia and to chronic activation and dysregulation of other bio-behavioral systems, including the hypothalamic-pituitary-adrenal axis (HPA), metabolic, and immune systems. Increased activation of the IFG was also found among individuals with ASD when they were required to identify face targets, despite lower accuracy in the task (Dichter et al., 2009). This was attributed to an attempt to compensate for an impairment in related cognitive processes due to cortical inefficiency. Similar hyper-activation of the OE was found in multiple studies in ASD (for a recent review refer to Chan & Han, 2020). Moreover, previous studies showed that lonely people have a reduction in fractional atrophy of white matter tracts linked to the IFG (Tian et al., 2014) as well as decreased white matter density in the IPL (Nakagawa et al., 2015), which may further reduce the effectiveness of the OE system. No differences were found in the dmPFC activity between high and low loneliness individuals. This may suggest that while the OE system is hyper-active, there may be an intact gap-monitoring system among lonely individuals. In essence, the results of this study suggest that lonely people may be exerting more neural effort in the OE system, however despite this they still achieve less optimal behavioral outcomes (Figure 8).

**Figure 8:**
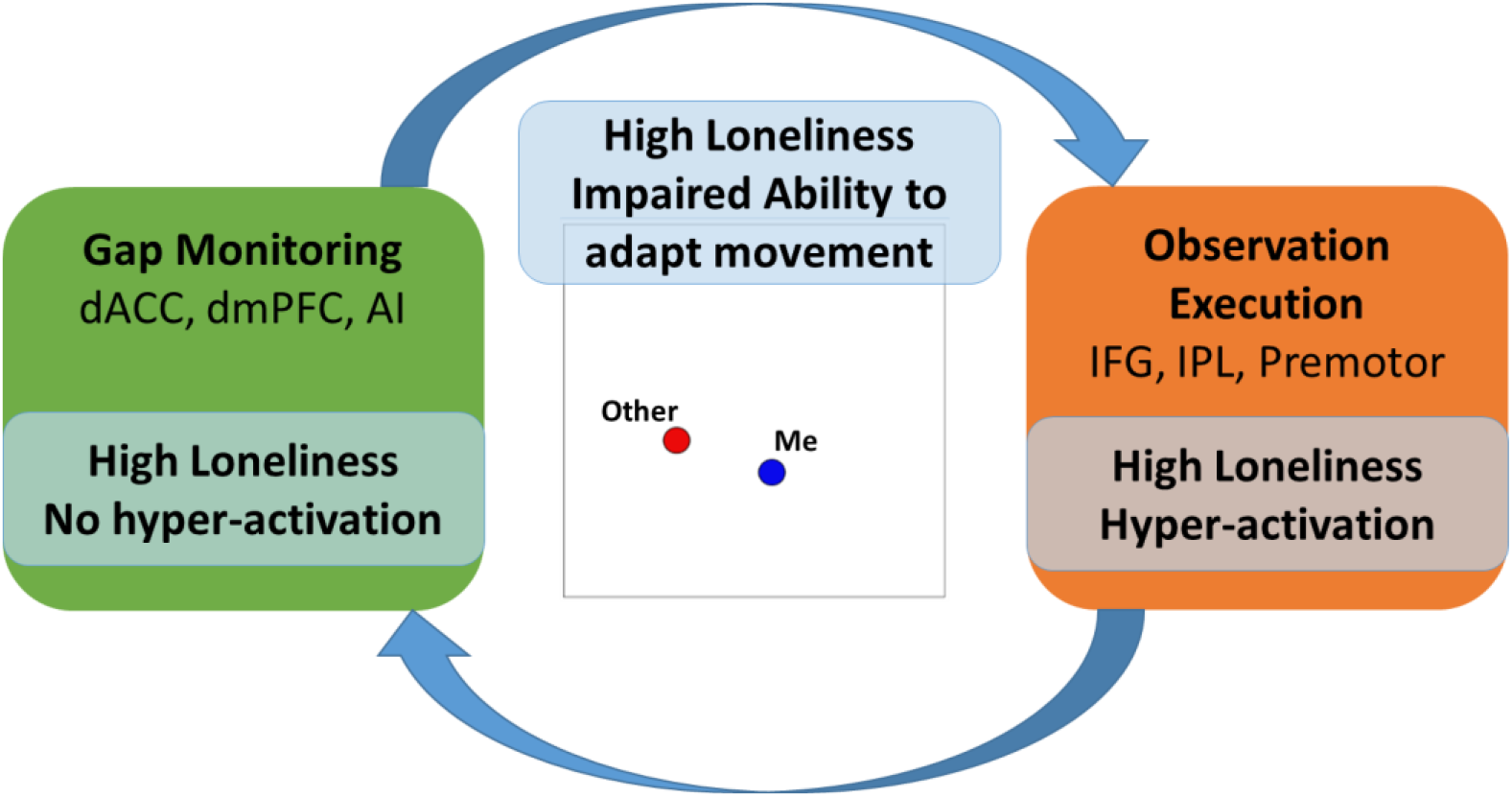
Proposed model of impaired synchronization in high loneliness individuals. The gap-monitoring system may be functioning properly, however there appears to be hyper-activation of the OE system, potentially to compensate for the impaired ability. Despite this hyperactivation, lonely individuals still experience difficulties in adapting their movement

Involvement of regions associated with the reward system (OFC, vmPFC, VS), which was also proposed in the interpersonal synchronization model (Shamay-Tsoory et al., 2019) to be important for the achievement of synchronization through signaling that optimal alignment was achieved, was not identified in this study. This may speak against the involvement of the reward system in movement synchronization, but it is also possible that longer periods of optimal alignment are required to stimulate the reward system. In addition, there is recent evidence that multiband sequences have lower power to detect reward-associated striatal activation (Srirangarajan et al., 2021).

As expected, participants’ enjoyment ratings were higher in the sync condition compared to the rand condition. However, participants also reported enjoying the free condition more than the rand condition. Therefore, it is possible that the increased enjoyment reported was not necessarily due to synchronization, but rather more related to the fact that participants knew they were interacting with a human. Moreover, despite previous accounts of lonely people reporting lower enjoyment from social interaction, in our study there were no significant differences in reported enjoyment between the high and low loneliness groups in any of the conditions. While other studies also failed to find a direct relation between enjoyment and loneliness (e.g., (Nezlek et al., 2002), it is also possible lonely individuals may have not enjoyed this interaction less since it was a virtual interaction, in which they typically feel more comfortable (Nowland et al., 2018). In addition, loneliness may be more related to other aspects of the experience, such as level of closeness or satisfaction from the interaction relationship (Mund et al., 2020) which were not measured in the current study.

During the free condition, we did not observe spontaneous synchronization, across both loneliness groups. Nonetheless, when examining the brain activity during the entire free condition and contrasting it with the control condition, it was apparent that brain regions relevant to synchronization were recruited, namely the IPL and the PFC. Therefore, it is possible that participants were recruiting the relevant regions but not to an extent that allows actual detectable synchronization. It is conceivable that the observed activations are related to the operation of the default mode network (DMN) (Buckner et al., 2008; Raichle et al., 2001), which is closely linked to social cognition (Mars et al., 2012; Smallwood et al., 2021). This might explain the difference between the free condition and the rand condition as participants knew that they are interacting with another person in the free condition. It is noteworthy that the activations in the PFC during the free condition were more lateral than in the sync condition, including the superior and dorsolateral parts of the PFC. The dlPFC was previously linked to approach-avoidance motivation conflict (Ironside et al., 2020; Rolle et al., 2021; Spielberg et al., 2012) and it is possible that this explains the activations during the free condition.

It should be noted that there were no differences in the zero-lag correlation score between the high and low loneliness groups. This can potentially be attributed to the fact that the zero-lag correlation is a dyad measurement, which does not reflect the individual contribution of each of the dyad members to the synchronization achieved. It is possible that high loneliness individuals contributed less to the synchronization, while their dyad partners compensated for their difficulties which resulted in intact dyadic performance. Future study designs can further test this possibility by specifically recruiting dyads in which both members have high loneliness versus dyads in which both members have low loneliness.

Reservations concerning the use of a naturalistic paradigm are warranted. While it has benefits in terms of validity and reliability, each participant interacted with a specific participant, and it may be claimed that their behavior could be different if they were to interact with different participants. To minimize the impact of this issue, we chose to focus on an individual measurement of contribution to the synchronization and the related neural activations and not the dyad measurement. That said, further research may be needed to confirm the findings of the study also in a more controlled setting. In addition, participants performed the task twice and it could be claimed that this would impact their neural and behavioral results. However, scanning order did not impact any of the behavioral or neuroimaging analyses and therefore it appears that this does not limit the ability to interpret the results.

In conclusion, we propose that lonely individuals may have an underlying impairment in interpersonal synchronization. We further propose that this is related to a hyper-activation of the OE system during synchronization, potentially as a compensation attempt for their impaired ability. Our study suggests that despite this hyper-activation, high loneliness individuals still achieve less optimal behavioral outcomes. Building on a model according to which all levels of alignment are related and involve the OE system (Shamay-Tsoory et al., 2019), we suggest that these difficulties may extend to emotional and cognitive synchronization as well. Interpersonal synchronization may therefore be a relevant area of intervention to ameliorate chronic loneliness, focusing on improving the lonely individual’s ability to individually contribute to synchronization. Targeting this may help improve the way lonely people experience social interactions and relationships, leading to increased sense of affinity, closeness, and satisfaction.

## Institutional Review Board Statement

The study was approved by the ethics committee of Tel Aviv University and the institutional review board at the Sheba Tel Hashomer medical center (approval number SMC-17-4132) and was conducted according to the guidelines of the Declaration of Helsinki

## Informed Consent Statement

Informed consent was obtained from all subjects involved in the study.

## 9 Captions to Figures

Figure 1: Illustration of the task design

Figure 2: Example of the zero-lag synchronization scores calculated over time for one dyad. The blocks of rand condition (left), free condition (middle) and sync condition (right) are presented. The × axis represents the time in seconds, and the Y axis is the synchronization score

Figure 3: High loneliness group had a lower following score in the sync condition. following score was higher in the sync condition, compared to free and rand condition. Error bars=95% confidence level (cl)

Figure 4: Brain activation in the sync condition>rand condition contrast, during the total run duration, across loneliness groups. Contrast thresholded at p<0.001 for the illustration. MNI coordinates of Axial/Coronal/sagittal view - (5, -40, 40).

Figure 5: Brain activation in the free condition>rand condition contrast, during the total run duration, across loneliness groups. Contrast thresholded at p<0.001 for the illustration. MNI coordinates of Axial/Coronal/sagittal view - (5, -40, 40)

Figure 6: Brain activation in the following periods sync>rand contrast, thresholded at p<0.001 for illustration. MNI coordinates of Axial/Coronal/sagittal view - (5, -40, 40).

Figure 7: high loneliness group presents a higher activation in the lIFG and rIPL during following periods. Error bars=95% confidence level (cl)

Figure 8: Proposed model of impaired synchronization in high loneliness individuals. The gap-monitoring system may be functioning properly, however there appears to be hyper-activation of the OE system, potentially to compensate for the impaired ability. Despite this hyperactivation, lonely individuals still experience difficulties in adapting their movement

